# Alpha-synuclein expression in neuron modulates the Japanese encephalitis virus infection

**DOI:** 10.1101/2023.09.25.559402

**Authors:** Anjali Gupta, Anshuman Mohapatra, Harpreet Kaur, Ajanta Sharma, Nitin Chaudhary, Sachin Kumar

**Affiliations:** Department of Biosciences and Bioengineering, Indian Institute of Technology Guwahati, Guwahati - 781 039, Assam, India; Division of Epidemiology and Communicable Diseases, Indian Council of Medical Research, Ansari Nagar, New Delhi - 110 029, India; Department of Microbiology, Gauhati Medical College, Guwahati – 781 032, Assam, India

**Keywords:** Japanese Encephalitis Virus, Alpha-synuclein, Antiviral, SOD1, Pathogenicity

## Abstract

Japanese encephalitis virus (JEV) stands as a prominent vector-borne zoonotic pathogen, displaying neurotropism and eliciting Parkinson’s disease (PD)-like symptoms among most symptomatic survivors. A characteristic feature of PD is aggregation of mutated α-synuclein (α-syn) that damages the dopaminergic neurons. Considering this link between JEV-induced PD-like symptoms and α-syn pathogenesis, we explored the role of α-syn in JEV infectivity in neuronal cells. Our investigation revealed significant increase in endogenous α-syn expression in JEV-infected cells. Additionally, treatment with exogenous α-syn (Exoα-syn) led to a substantial reduction in JEV replication, suggesting its anti-JEV effect. Furthermore, Exoα-syn treatment led to the upregulation of superoxide dismutase 1 (SOD1) and reduction in reactive oxygen species (ROS). The results were validated by endogenous α-syn-silencing, that decreased SOD1 level and raised ROS level in neuronal cells. Similarly, the SOD1 inhibition via LCS-1 also intensified ROS and JEV infection. Overall, our results suggest that α-syn exerts an anti-JEV effect by regulating protein involved in oxidative stress inside neuronal cells. This study contributes valuable insights into the interplay between α-syn expression and JEV infectivity, shedding light on avenues to further investigate the potential role of α-syn in JEV pathogenesis.

**Importance:** Japanese encephalitis virus (JEV) poses significant threat particularly to children. Despite extensive research efforts, the development of effective treatments against JEV has been impeded. One of the major setbacks is a lack of comprehensive understanding of neurotropism. The study focuses on alpha synuclein (α-syn), a neuronal protein, and aims to determine its role in JEV pathogenesis. Present study reveals that host cell upregulates α-syn in response of JEV infection. α-syn restrains JEV propagation by modulating superoxide dismutase 1 (SOD1) expression which further blocks JEV-induced ROS generation. Endogenous α-syn silencing led to decrease in SOD1 expression and increased viral titer. α-syn plays crucial role in counteracting oxidative stress through SOD1, which is essential for limiting JEV replication. This study provides broader implications for antiviral strategies and its possible role in neurodegenerative diseases; however, there is still much to explore, particularly regarding its aggregation kinetics in JEV infection.

## Introduction

Japanese encephalitis (JE) is a serious infectious disease of central nervous system. Approximately 15% of JE patients die during the acute phase of illness, and most survivors present with neurological aftereffects, including striatal dysfunction (1, 2). JEV replication has been found to be localized mainly at thalamus, substantia nigra, basal ganglia, cerebral cortex, cerebellum, and brain stem in clinically infected patients (3, 4). Neurological complications associated with the critical phase of illness include encephalopathy and movement disorders similar to parkinsonism (5, 6). Parkinsonism include symptoms like tremors, a masked face, microphonia, impaired consciousness, saccadic eye tracking, and neck stiffness (7, 8). Apart from JEV, these symptoms are associated with a variety of other enveloped RNA virus infections like West Nile virus (WNV) and Dengue virus (DNV) (9, 10). Neuron specific proteins are likely to play a major role in JEV prognosis. Recent work has shown that innate immune responses to viral infections in the CNS contribute to neuron-specific injury patterns (11). However, expression of a neuron-specific restriction factor for viral infections has not been well studied.

Alpha synuclein (α-syn) is one of the neuron-specific proteins regulating functions associated with the synapse, such as synaptic plasticity, neurotransmitter release, dopamine metabolism, and vesicle trafficking (12). It is extensively expressed in the thalamus, substanti nigra, and basal ganglia. It is a protein of 140 amino acid residues with an amphipathic alpha-helix structure similar to apolipoproteins at its N-terminal sequence due to seven 11-mer repeats with a KTKEGV consensus sequence (13). This N-terminal domain imparts lipid binding propensity to the protein (14, 15). The C-terminal region of α-syn (residues 96–140) is highly acidic and largely unstructured. α-syn has been reported in the literature to cause membrane remodeling by inducing membrane curvature, converting large vesicles into highly curved membrane tubules, micelles and vesicles (16–18). It also inhibits phospholipases D1 and D2, and is involved in the cleavage of membrane lipids (19, 20). Its C-terminal part is known to be the target of various post-translational modifications like phosphorylation, ubiquitination, and SUMOylation. These modifications are believed to be responsible for its altered binding affinities or interactions with cellular proteins and lipids (21). The role of α-syn has largely been explored in the context of Parkinson’s disease (PD). In most of the Parkinson clinical patients, missense mutations in the α-syn gene (A30P, E46K, A53T) are likely to cause its aggregation and formation of Lewy bodies (LBs) in the cytoplasm (22). These aggregated α-syn induce toxicity by increasing reactive oxygen species (ROS) inside neurons, eventually causing damage to mitochondria and other organelles (23, 24). In a biological system, ROS, such as hydrogen peroxide (H_2_O_2_), superoxide anion (O_2_•−), hydroxyl radical (•OH), and singlet oxygen (^1^O_2_) are oxidants and mediators of cell injury, disease, homeostasis, and signaling activation (25, 26). Moreover, the function of native or monomeric α-syn is not studied in as much detail as that of mutants. Contrary to PD, α-syn plays a neuroprotective role and is also important in pathogen-activated immune responses and lymphocyte maturation (27). In microglia cells, native α-syn leads to an increase in Th1 and Th2 cytokine expressions (28). Moreover, α-syn has been reported to directly interact with supercoiled DNA and RNA-interacting proteins to regulate specific gene expression (29). The nuclear translocation of α-syn is found to be important in modulating DNA repair (30). Taken together, these studies suggest that α-syn could play a pivotal role in pathophysiological responses involved in inflammatory disease to invading pathogens.

Even though there is extensive literature on α-syn, its role in viral pathogenesis has not been studied well. Recent studies showed that JEV exploits dopamine signaling and modulates its level to facilitate viral infection. In addition, dopamine D2 receptor (D2R) helps JEV entry by activating phospholipase C (PLC) signaling cascade (31). Both dopamine level and D2R activity depend on α-syn, and therefore, are of critical importance to maintain synaptic homeostasis (32).

The purpose of this study is to explore multifaceted potential of α-syn in JEV pathogenesis. To achieve the objectives, mouse neuronal cells were studied upon JEV infection, endogenous α-syn silencing, with α-syn overexpression, and upon treatment with exogenous α-syn (Exoα-syn).

## Material and Methods

### Cells and Virus

The mouse neuronal cells (Neuro2a) and baby hamster kidney cells (BHK-21) were procured from National Centre for Cell Science, Pune, India. The JEV stock preparation, along with all the plaque assays, were carried out in BHK-21. Both BHK-21 and neuro2a cells were maintained in Dulbecco’s modified eagle medium (DMEM), containing 1X penicillin streptomycin antibiotic with 10% fetal bovine serum (FBS) in 5% CO_2_ at 37 °C. The JEV strain SA14-14-2 (GenBank accession number JN604986) was obtained from the Department of Health and Family Welfare, Government of Assam, India. For stock preparation, BHK-21 cells at 80% confluency were infected with JEV at MOI 0.1. After 2 h of virus adsorption, infection media was removed and replaced with DMEM media containing 2% FBS. The infected cells were lysed by multiple freeze thaw cycles and clear lysate was collected 72 h post-infection and stored at -80 °C.

### Expression and purification of exogenous α-syn protein

Exoα-syn protein was expressed in *E. coli* BL21(DE3) expression system. The pET21a-alpha-synuclein was obtained from Michael J Fox Foundation MJFF (Addgene plasmid # 51486) as a kind gift. Briefly, the plasmid was transformed into *E. coli* BL21(DE3) cells. The recombinant cells were grown up to an OD_600_ = 0.6 in Luria-Bertani broth enriched with 100 μg/ml ampicillin at 37 °C and 150 rpm shaking. The culture was induced using 1 mM IPTG and further grown for 15 h at 25 °C. The cell suspension was pelleted at 7000 rpm for 5 min and resuspended in 100 ml Buffer A (20 mM Tris-Cl, 5 mM EDTA, pH 8.0) containing 1 mM PMSF. The cells were sonicated for 45 min with 8 s ‘ON’ and 22 s ‘OFF’ cycle at 33% amplitude. The sonicated sample was boiled at 95 °C. The insoluble white precipitate was then pelleted at 20,000 g for 1 h. The clear supernatant was passed through a 0.45 μm filter before further purifying using chromatographic methods as described elsewhere (33).

### Characterization of exogenous α-syn protein

The purified protein was characterized using SDS-PAGE. The molecular weight of the purified protein was determined using MALDI-TOF on a Bruker Autoflex Speed MALDI-TOF/TOF mass spectrometer, USA using α-cyano-4-hydroxycinnamic acid matrix. The dynamic light scattering (DLS) measurements of the freshly purified Exoα-syn were performed using a Malvern Zetasizer NANO-ZS DLS, UK instrument. The DLS measurements were performed at room temperature using a 173° backscattering angle and analyzed using the Zetasizer software version 7.11.

## Time-course analysis of α-syn expression

Neuro2a cells were seeded in a 6-well plate at a density of 2 × 10^6^ cells/well in DMEM supplemented with 10% FBS. After 12 h incubation, the cells were washed with PBS, followed by addition of 500 µl diluted JEV stock at 0.1 MOI per well. The cells were collected post 12-96 h of incubation and processed for RNA isolation and protein lysate preparation.

### *In vitro* anti-JEV activity

Neuro2a cells were seeded in 6-well plate at a density of 2 × 10^6^ cells/well. For co-treatment assay, JEV at 0.01 MOI was incubated with three different concentrations of Exoα-syn (0.25, 0.5 and 1 µM) for 30 mins along with appropriate controls. Following incubation, the mixture was added on top of cells and kept for 2 h for adsorption. The medium was removed post-infection, and fresh 2 ml DMEM with 2% FBS was added in each well.

For post-treatment assay, Exoα-syn was added to the cells after 2 h of JEV adsorption, whereas for pre-treatment assay, Exoα-syn was added 6 h before JEV infection. Cells and supernatant from differentially treated (CON, JEV, JEV along with 0.25, 0.5, 1 µM of Exoα-syn) samples were collected at 72 h time point. JEV genomic RNA and its proteins were detected by real-time PCR and western blot analysis, respectively. In addition, JEV titration was done by plaque assay.

### Small Unilamellar Vesicles (SUVs) preparation

1-palmitoyl-2-oleoyl-*sn*-glycero-3-phosphocholine (POPC), 1-palmitoyl-2-oleoyl-*sn*-glycero-3-phospho-*L*-serine sodium salt (POPS), 1-palmitoyl-2-oleoyl-*sn*-glycero-3-phosphoethanolamine (POPE), and sphingomyelin (SM) were precured from Avanti Polar Lipids, Inc, UK. The lipids POPC:POPS:POPE:SM in 57:25:3:15 molar ratio were aliquoted into a clean glass tube. As described elsewhere, a thin lipid film was prepared by slowly swirling the lipids in the glass tube under a stream of nitrogen gas (34). The lipid thin film was left to dry overnight in a vacuum desiccator. The overnight-dried lipid film was hydrated using 1 ml of phosphate buffer (PB) for six hours. The hydrated lipid was subjected to vigorous vortexing and five freeze-thaw cycles in an ice bucket and warm water bath. The lipid suspension was allowed to reach room temperature and sonicated in a water bath sonicator until the solution cleared out to form SUVs. The SUVs prepared were analyzed using DLS.

### Circular Dichroism (CD) analysis

Circular dichroism spectroscopic measurements were carried out on a J-1500 Jasco CD spectropolarimeter in 25 mM phosphate buffer, pH 7.5. The spectra were recorded at 2 μM protein concentration in the absence of lipids and in the presence of SUVs at 1:200 protein:lipid ratio. Each spectrum was recorded with eight accumulations from 300 nm up to 190 nm with 0.1 nm data pitch, 1 nm bandwidth, 2 sec D.I.T., and a 100 nm/min scan speed. The CD spectra were smoothed 25 points using the second polynomial function of the Savitzky-Golay algorithm.

### Calcein-loaded vesicle preparation

The lipid film with same lipid composition *i.e.* POPC:POPS:POPE:SM (57:25:3:15) was prepared in a clean glass tube and hydrated using 80 mM calcein solution in PB, pH 7.5 The calcein-loaded SUVs were prepared by sonication as described above and cleared through a HiTrap PD10 desalting column in phosphate buffer. The early fractions containing calcein-loaded SUVs were collected and characterized with DLS, and used for calcein release assay. The fluorescence emission was recorded for 600 s in a Jasco FP-8500 spectrofluorometer (35). The protein lipid-interaction studies were carried out with 0.5 μM Exoα-syn at 1:10, 1:25 and 1:50 protein:lipid molar ratios. Triton X-100 (1%v/v) was used a positive control.

### Overexpression and silencing of α-syn

Plasmid pHM6-alphasynuclein-WT, a kind gift from David Rubinsztein (Addgene plasmid # 40824), was used for the overexpression of α-syn in neuro2a cells (36). The neuro2a cells were seeded in four 35 mm cell culture petri dishes at a density of 2 × 10^6^ cells/ dish. The two dishes were transfected with 2 µg of plasmid using Lipofectamine 3000 (Invitrogen, USA) following the manufacturer’s protocol. After 12 h, a transfected dish and an untransfected one were infected with 0.1 MOI JEV. After 72 h of incubation, supernatant and whole cell lysate were collected for plaque assay and protein estimation.

The silencing of endogenous α-syn was carried out *via* combined transfection of two siRNAs at 75 pmol total concentration. The following sequences of siRNAs sense strands, targeting two different exons of α-syn, were used: CUAAGUGACUACCACUUAU[dT][dT] and CACAGGAAGGAAUCCUGGA[dT][dT] (Merck, Germany). The transfection reagent Lipofectamine RNAiMAX (Invitrogen, USA) was used following manufacturer’s protocol, along with siRNA universal negative control #1 (SIC001, Sigma-Aldrich, Germany). The knockdown of α-syn was estimated after 24 h of transfection by real-time PCR analysis and western blotting.

### Gene expression and viral particle estimation

Total RNA was isolated using RNAiso reagent (Takara, Japan) and converted into cDNA using iScript™ cDNA Synthesis Kit (Biorad, USA). Real-time PCR analysis was carried out in system QuantStudio5 (Applied Biosystems, USA) using Power-up SYBR master mix (Invitrogen, USA). GAPDH was used as an internal control and values are represented as mean fold change with respect to cell control (CON). Protein estimation were carried out using SDS-PAGE and western blotting. Western blotting were performed using antibodies specific to JEV nonstructural protein 1 (NS1) (GTX633820, GeneTex, USA), α-syn (32-8100, Invitrogen, USA), SOD1 (A0274, Abclonal, USA), and DJ-1(5933), SQSTM1 (39749), and NQO1 (62262) from CST, USA. The β-actin (MA1-91399, Invitrogen, USA) or GAPDH (MA1-16757, Invitrogen, USA) were used as a loading control for all the experiments. The relative quantification was done using ImageJ software. For JEV particle quantification, plaque assay was performed in BHK-21 cells using standard protocol (37).

### ROS estimation

For all flow cytometry studies, neuro2a cells were seeded at 2 × 10^6^ cells/well density in 6-well plate and incubated for 12 h at 37 °C, 5% CO_2_ incubator. ROS was estimated in JEV-infected and mock-infected cells at 24, 48, 72, and 96 h time points. Briefly, 0.1 MOI JEV was added in 4 wells and equal number of wells were kept as controls. The cells were detached every 24 h post-infection, resuspended in 500 μl PBS containing 10 μM of 2′,7′-dichlorodihydrofluorescin diacetate (DCFH-DA) dye, and incubated for 30 min at 37 °C. The stained cells were analyzed using flow cytometry (BC, CytoFlex S Analyser). FITC filter channel was used to record the emission of DCFH-DA dye. Flow cytometry analysis was done post-treatment with compounds; H_2_O_2_, ascorbic acid, Exoα-syn, and LCS-1 at different time intervals and concentrations. Ascorbic acid and Exoα-syn treatment were given 12 h before flow cytometry at 0.2 and 1 μM concentrations, respectively. Cells were treated with 500 μM of H_2_O_2_ for 6 h, whereas for LCS-1, ROS was estimated after 6 and 12 h of treatment with 5 μM of LCS-1.

## Statistical analysis

The data presented in the results are the average of three independent experiments. Significance values were calculated through two-tailed analyses using Student’s t-test in Microsoft Excel. Asterisk marks; ** represent a p-value < 0.01, while * represents a p-value < 0.05.

## Results

### Modulation of α-syn post-JEV infection in neuronal cells

Time course analysis of α-syn expression upon JEV infection was carried out in neuro2a cells for 96 h. The mRNA estimation using real-time PCR indicated gradual increase in α-syn from 24 h till 60^th^ h. The maximum ∼3.7-fold change, with respect to the mock-infected neuro2a cells, was obtained at 60^th^ h. Thereafter, with the rapid increase of JEV E mRNA, α-syn mRNA decreased to ∼2.4-fold level (Figure 1A and 1B). At protein level, gradual increase in α-syn was observed with time in both mock-infected and JEV-infected cells. However, with respect to mock-infected neuro2a, there was significant upregulation of α-syn in JEV-infected cells at all the time points (Figure 1C).

**Figure 1.**
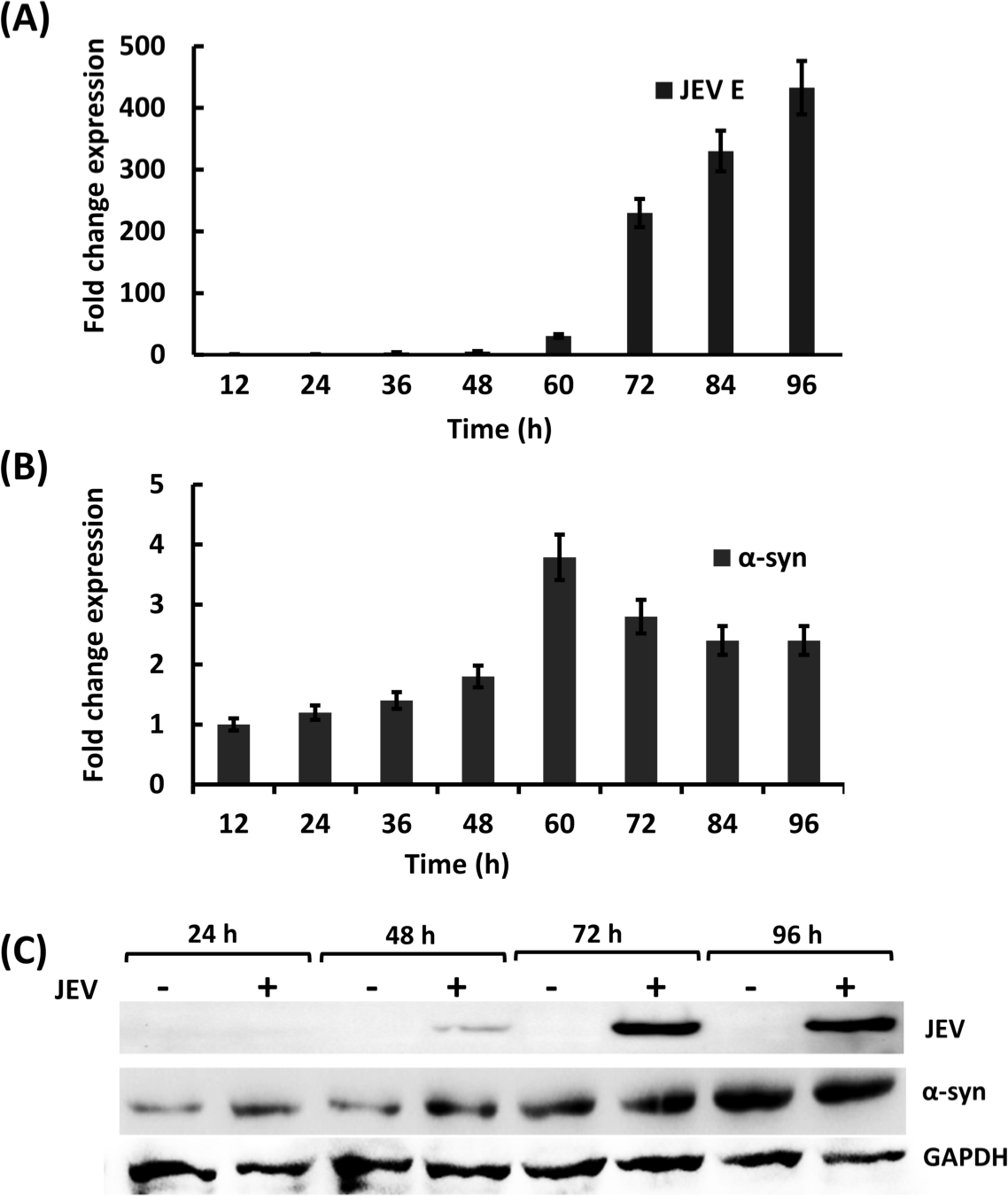
Quantification of endogenous α-syn in neuro2a cells. Real-time PCR analysis of viral mRNA at 12 h interval till 96 h post-infected with 0.1 MOI JEV (A). Fold change expression of α-syn in JEV-infected neuro2a cells compared to mock-infected cells till 96 h where GAPDH is used as a control (B). Fold change is calculated keeping mock-infected cells as control. Western blot analysis of α-syn protein in JEV-infected neuro2a cells compared to mock-infected cells till 96 h (C).

### α-syn inhibits JEV replication

The purified Exoα-syn was characterized before proceeding towards experiments. The anion exchange-purified chromatogram of Exoα-syn showed elution around 55 min on a gel filtration column (Supplementary Figure 1A). The different 55 min fractions were run on 12% SDS PAGE that showed ∼15 kDa protein band (Supplementary Figure 1B). The MALDI-TOF mass spectrum showed peaks at 7233.25 and 14,476.18 m/z values (Supplementary Figure 1C). The peak at 14,476.18 corresponds to the [M + Na]^+^ adduct of Exoα-syn whereas the smaller m/z value corresponds to the doubly-charged Exoα-syn. DLS analysis revealed a single peak with mean hydrodynamic diameter of 8.01 nm with PDI of 0.246, indicating a monodispersed protein preparation (Supplementary Figure 1D). This hydrodynamic diameter is comparable to those reported in the literature for monomeric α-syn (38, 39). To analyze the role of Exoα-syn in JEV replication, differential treatment of neuro2a cells was carried out with Exoα-syn along with JEV infection. Significant downregulation of NS1-specific mRNA was observed in pre- and post-α-syn treatment. In pre-treatment, the viral mRNA gradually decreased with an increase in α-syn concentration (Figure 2A). Treatment with 1 µM α-syn caused around 80% reduction in the mRNA level. Similar results were obtained for the post-treatment samples as well. Around 75% reduction in the mRNA level was observed at 1 µM α-syn concentration. Inhibition in pre-treatment was slightly better than post-treatment. In co-treatment, no significant difference in viral mRNA levels was observed at any Exoα-syn concentration. The cellular protein estimation indicated the similar trend as observed in gene expression studies. Upon α-syn pre-treatment (0.25 µM α-syn), JEV protein amount reduced to about 84% compared to untreated JEV infected control.

**Figure 2.**
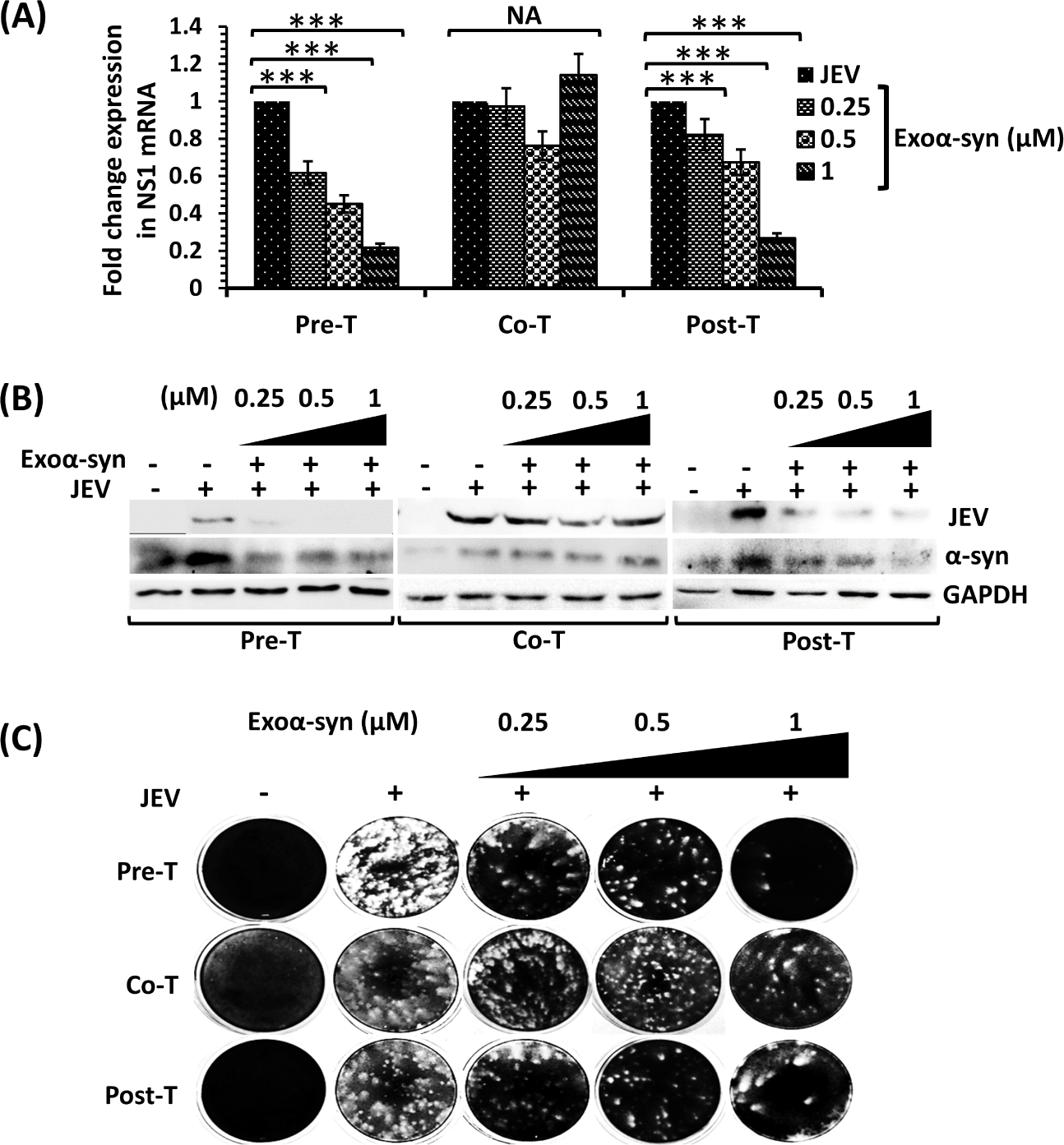
Modulation of JEV in neuro2a cells after co-treatment, pre-treatment, and post-treatment of Exoα-syn at 0.25, 0.5, and 1 µM concentrations. JEV mRNA quantification after 72 h of infection in differentially-treated cells. Fold change is calculated with respect to uninfected cells as control (A). Western blot analysis to quantify the viral protein after 72 h of JEV infection in cells differentially treated with Exoα-syn (B). Virus titration of supernatant collected after 72 h of JEV-infected cells differentially-treated with Exoα-syn protein (C).

In post-treatment samples, JEV protein amount got reduced to 69-91% compared to infected control. Endogenous α-syn expression also seems to get regulated with JEV replication. The JEV-infected control exhibited the highest α-syn expression, whereas cells with lower JEV replication displayed reduced α-syn levels both in pre- and post-treatment samples (Figure 2B). In co-treatment, uniform JEV replication across differentially treated cells resulted in consistent endogenous α-syn expression. In plaque assay, more than 82 % decrease in JEV titer were observed in 0.25 µM of Exoα-syn pre- and post-treated cells. In co-treatment, about 32% decrease in JEV titer was observed only at 1 µM of Exoα-syn treatment (Figure 2C).

As α-syn is known to bind lipid vesicles, its interaction with SUVs made up of lipids that constitute viral lipid bilayer was investigated. The diameters of plain and calcein-loaded SUVs, as determined by DLS, were around 32 nm and 55 nm, respectively (Supplementary Figure 2A). Circular dichroism spectrum of Exoα-syn in the absence of SUVs suggest a random coil conformation, as is expected for a monomeric α-syn. In the presence of SUVs, on the other hand, the protein folds to take up an α-helical conformation (Supplementary Figure 2B). Disruption of lipid vesicles upon Exoα-syn binding, if any, was investigated through calcein release assay. Calcein was loaded in the SUVs at self-quenching concentration. An increase in fluorescence intensity upon α-syn addition would imply membrane disruption. The extent of fluorescence is directly proportional to the extent of vesicle disruption. Treatment of calcein-loaded SUVs with Exoα-syn did not cause any appreciable enhancement in calcein fluorescence (Supplementary Figure 2C) suggesting that binding of Exoα-syn to SUVs does not disrupt them. The results obtained were in consonance with *in vitro* co-treatment study.

### Effect of overexpression and silencing of α-syn on JEV replication

The overexpression study was carried out by transfecting neuro2a cells with plasmids encoding wild type α-syn and GFP, separately. Cellular protein analysis revealed about 68% reduction in JEV protein in α-syn overexpressing cells than GFP control. Concurrently, about 61% lower JEV titer was observed in α-syn overexpressing cells than JEV infected cells expressing GFP (Figure 3A). Further, the silencing of endogenous α-syn was carried out through transfection of α-syn specific cocktail of two siRNAs. The downregulation was estimated using real time PCR and by western blot analyses. The endogenous α-syn mRNA was decreased by 57% and 64% compared to untransfected and -ve siRNA transfected neuro2a cells, respectively. A significant difference was observed at protein level as well, where α-syn-specific siRNA transfection downregulated α-syn by more than 76% as estimated by ImageJ software (Figure 3B). To estimate the replication of JEV upon α-syn silencing, cells were infected at 0.1 MOI after 12 h of transfection. Beside absence of any significant difference in viral protein, the JEV titer from the supernatant of α-syn-silenced cells showed up to 37% and 64% increase as compared to untransfected, and -ve siRNA-transfected JEV-infected cells, respectively (Figure 3C).

**Figure 3.**
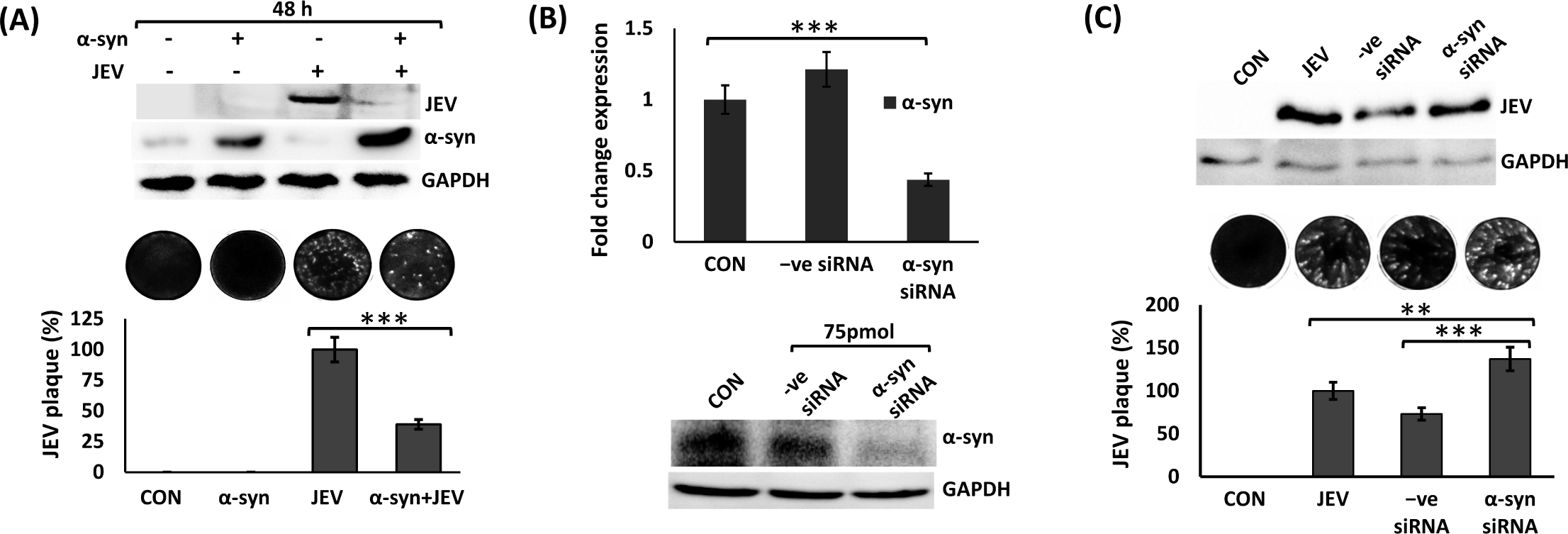
Modulation of JEV upon up and down regulations of α-syn gene. Western blot analysis of JEV protein after 48 h of its infection in cells transfected with plasmid expressing α-syn (A). The graph representing the percentage reduction in the plaque number. Downregulation of α-syn mRNA by siRNA treatment in neuro2a cells (B). Western blot analysis of JEV protein and plaque assay of neuronal cells transfected with siRNA and infected with JEV at 0.1 MOI (C).

### α-syn modulates oxidative stress via decreasing ROS in neuro2a cells

We investigated the ROS levels upon JEV infection, and upon treatment with ascorbic acid, and H_2_O_2_, two oxidative stress modulators, and Exoα-syn protein. JEV was infected at 0.1 MOI in neuro2a cells and ROS was estimated using DCFH-DA dye using flow cytometry. ROS during early stage of infection was comparable to mock-infected neuro2a cells. Their level, however, increased by about 59% in JEV-infected cells 72 h post-infection (Figure 4A). The H_2_O_2_ and ascorbic acid are positive and negative modulators of oxidative stress, respectively. As revealed by flow cytometric analysis, treatment with H_2_O_2_ resulted in ∼37% more ROS compared to untreated control cells after 6 h of treatment. Ascorbic acid and Exoα-syn both decreased ROS by about 19% and 16%, respectively compared to untreated control after 12 h of treatment (Figure 4B). These data revealed that Exoα-syn reduces intracellular ROS. Further, experiments were conducted to understand if JEV replication gets influenced by increase or decrease in ROS levels. We observed about 53% increase in JEV viral particle in the supernatant of H_2_O_2_ treated cells compared to untreated JEV-infected cells. Similar to Exoα-syn, ascorbic acid also negatively regulated JEV replication by approximately 55% compared to JEV control group (Figure 4C).

**Figure 4.**
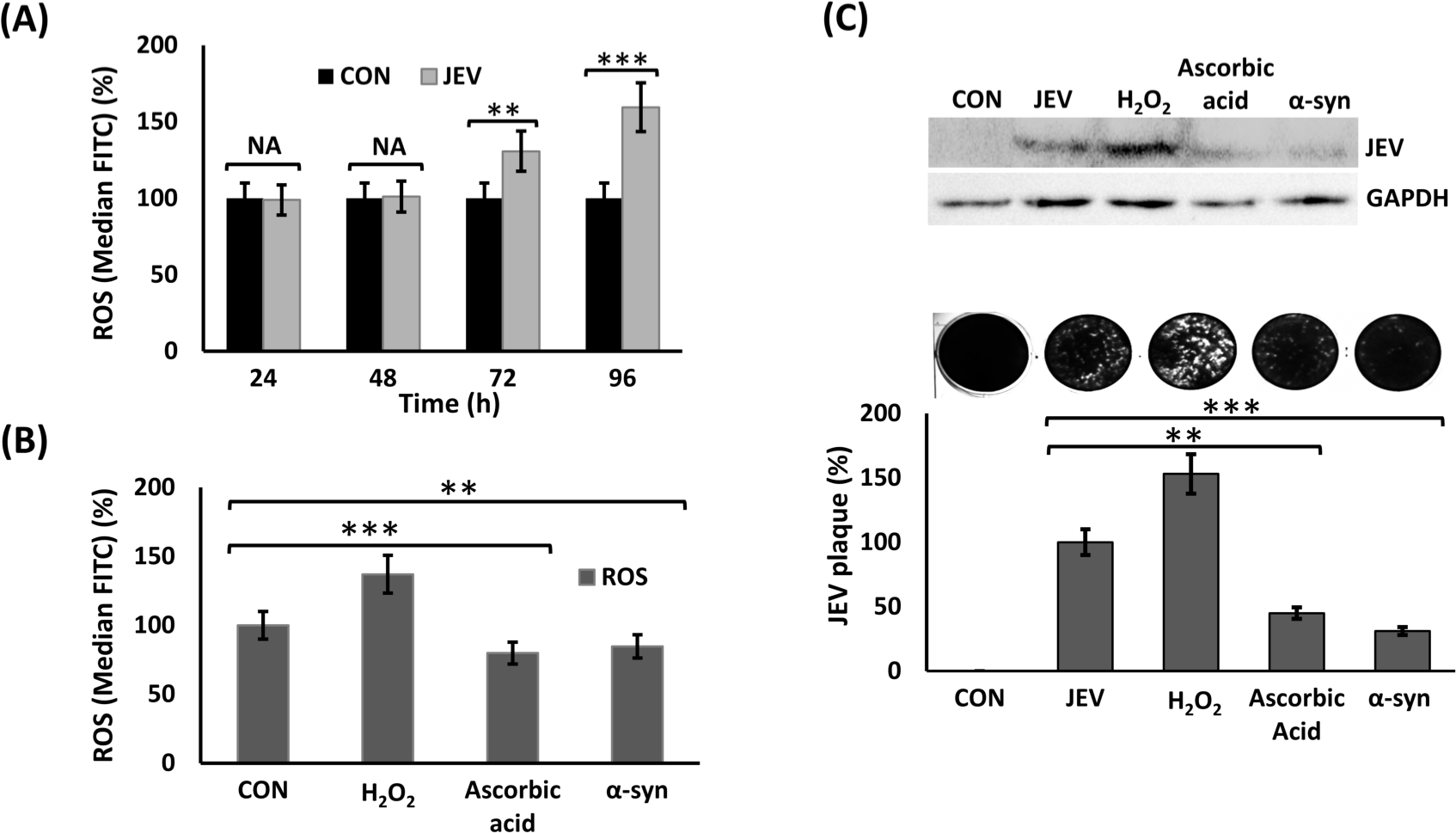
Reactive oxygen species (ROS) estimation using flow cytometry. Flow cytometry analysis to quantify ROS in JEV infection at 24 h interval till 96 h. DCFH-DA dye was used to stain the cells 30 min prior estimation in live cells in FITC channel. Graph representing median FITC (%) in mock-infected and JEV-infected cells (A). Median FITC (%) of ROS estimation in cells 6 h post-treatment with H_2_O_2_, 12 h post-ascorbic acid and α-syn protein treatment and untreated control (B). Modulation of JEV infection post-treatment with H_2_O_2_, ascorbic acid, α-syn protein, and untreated control. Western blot analysis along with viral titration through plaque assay from the whole cell lysate and supernatant of the treated cells, respectively (C). Graph representing the percent plaque numbers in differentially treated cells.

### α-syn positively regulates SOD1 expression

Neuro2a cells were treated with Exoα-syn at 0.25, 0.5, and 1 µM concentrations for 24 h, and cellular protein analyzed using western blotting. A significant increase in SOD1 level from ∼2.2 to ∼4.7-fold compared to untreated cells was observed with increasing concentration of Exoα-syn. No appreciable difference in SQSTM1, DJ-1, and NQO1 in either of the Exoα-syn concentrations was observed (Figure 5A). To validate whether SOD1 is modulated via endogenous α-syn, the SOD1 expression post α-syn silencing was analyzed. SOD1-specific mRNA was reduced to about 37% and 56% compared to -ve siRNA-treated cells and untreated control cells, respectively. At protein level, SOD1 protein was upregulated by about 17% in -ve siRNA-treated cells compared to untreated cells. The SOD1 downregulation in α-syn knockdown cells was found to be reduced by around 59% and 42% compared to -ve siRNA-treated and untreated cells, respectively (Figure 5B).

**Figure 5.**
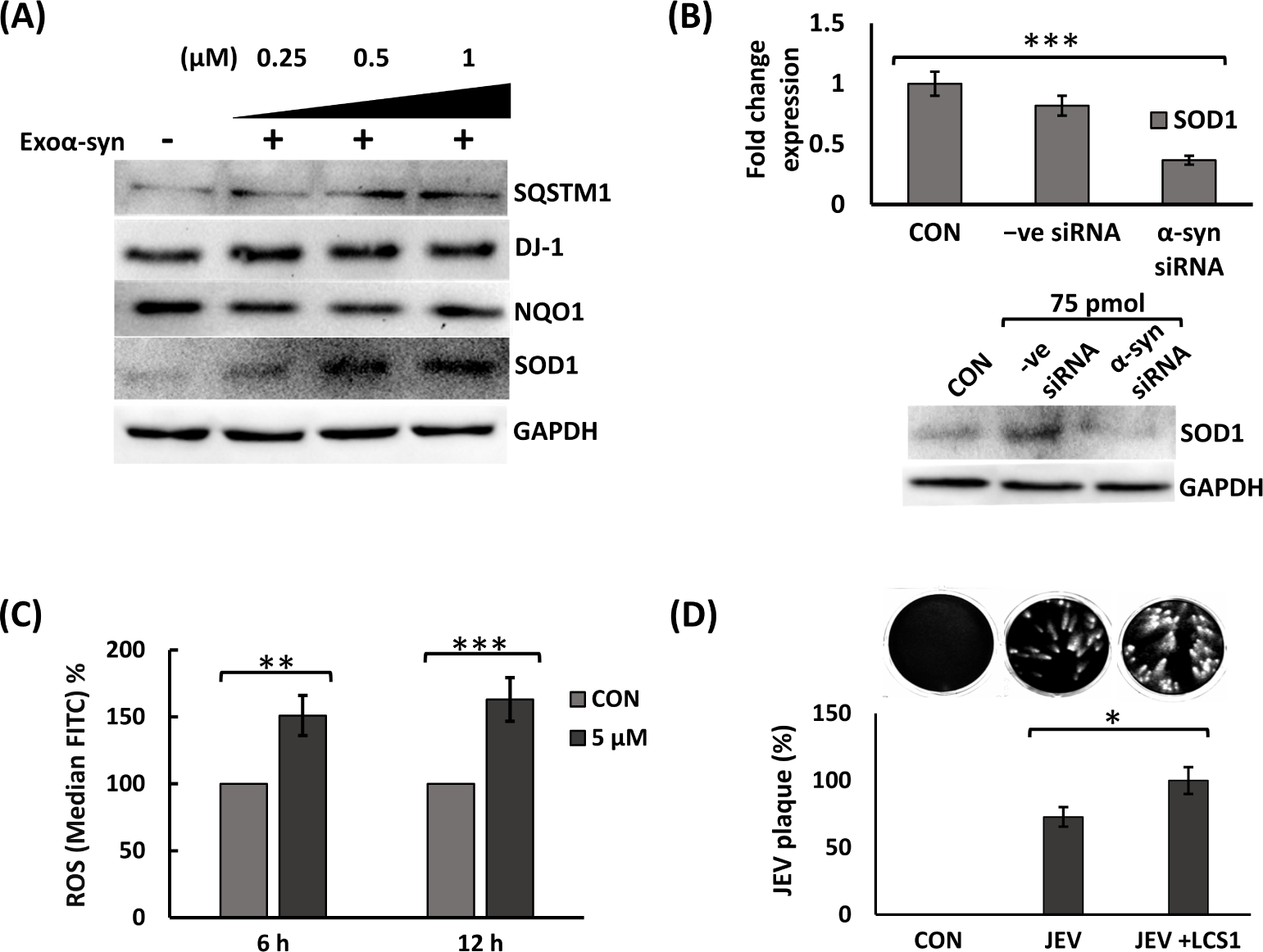
Modulation of genes involved in oxidative stress post-treatment with Exoα-syn protein at 0.25, 0.5, and 1 µM concentrations. Western blot analyses of SOD1, SQSTM1, DJ-1, and NQO1 proteins after 24 h of treatment. Untreated neuro2a cells are used as control cells and GAPDH as loading control (A). Modulation of SOD1 post-α-syn gene silencing. Real-time PCR and western blot analyses to estimate SOD1 specific mRNA and protein post α-syn siRNA treatment (B). Fold change is calculated by keeping SOD1 expression in untransfected (CON) cells as baseline. Modulation of ROS post-treatment with SOD1 inhibitor LCS-1. Graph representing median FITC (%) post 6 h and 12 h of 5 µM LCS-1 treatment in neuro2a cells (C). Untreated (CON) neuro2a cells are kept as control cells. Modulation of JEV post-treatment with 5 µM LCS-1. Viral titer analyses through plaque assay and graph representing plaques (%) in differentially-treated cells (D).

### Increase in JEV infectivity post-treatment with LCS-1

To check the role of SOD1 in JEV infection, the neuro2a cells were treated with LCS-1, a known inhibitor of SOD1 activity. Intracellular ROS level was increased by ∼51% and ∼63% compared to untreated neuro2a cells after 6 h and 12 h treatment of 5 µM LCS-1, respectively (Figure 5C). Moreover, it also enhanced the JEV replication by ∼23% post 12 h LCS-1 treatment (Figure 5D).

## Discussion

Neurotropic viruses, mainly JEV, pose a significant threat to children, leading to encephalitis-related fatalities. Despite intensive research endeavors, the development of effective treatments against these viruses has been impeded by the challenges of breaching the blood-brain barrier and a lack of comprehensive understanding of neurotropism (40, 41). To bridge these gaps and enhance therapeutic approaches, we investigated the function of a specific neuronal protein, α-syn in JEV pathogenesis. α-syn has been extensively studied in the context of PD and is found to be the primary reason of PD associated death of dopaminergic neurons. Recently, many clinical studies have uncovered the resemblance of Parkinson-like symptoms in JEV-infected patients. This prompted us to explore whether α-syn contributes to promoting JEV progression or inhibiting its replication.

We performed time-course expression analyses of α-syn in JEV-infected and mock-infected neuro2a cells. The results revealed a marked upregulation of endogenous α-syn at both mRNA and protein levels during the late phase of JEV replication in neuronal cells. This upregulation of α-syn is consistent with previous reports of increased α-syn levels in the brains of patients infected with other viruses, such as WNV and human immunodeficiency virus (HIV) (42, 43). To investigate the potential antiviral role of α-syn, we conducted experiments involving wild-type Exoα-syn. Pre- and post-treatment with Exoα-syn exhibited resistance to JEV infection. These findings align with previous studies that showed α-syn’s antiviral properties against WNV, wherein the brains of α-syn-knockout mice exhibited increased infectious viral particles and mortality compared to control mice (42). Intriguingly, our co-treatment experiments, wherein α-syn was administered along with JEV, did not show a significant antiviral effect. We hypothesized that Exoα-syn exerted its antiviral function through modulation of host cellular mechanisms rather than direct interference with the physiochemical properties of JEV virion particles or the viral entry process. We explored this hypothesis via investigating the biophysical interaction between Exoα-syn and calcein dye-encapsulated small unilamellar vesicles (SUVs) designed to mimic the lipid membrane of JEV (44). Using circular dichroism spectroscopy and calcein release assay, we found that Exoα-syn underwent a conformational change from random coil to α-helical conformation upon interaction with SUVs. Remarkably, this interaction did not cause any disruption in the lipid SUVs, as no calcein release was observed. While α-syn is known to exhibit lipid binding affinity under cellular conditions, our results suggest that it does not significantly disrupt the viral lipid membrane or inhibit viral entry during JEV infection.

The behavior or function of α-syn seems to differ based on cell type, its mutational status, and the conformation of the protein (aggregated or non-aggregated). Several studies have demonstrated a relationship between mutated and aggregated α-syn and oxidative stress, leading to neurotoxicity and its involvement in neurodegenerative diseases (45–47). However, besides the well-documented neurodegenerative activity of α-syn, the role of monomeric and wild-type α-syn has been found to be neuroprotective in many studies (48, 49). The toxicity of wild-type α-syn is specifically dependent on the presence of dopamine. In the absence of dopamine, in human cortical neurons, α-syn protects the cells and significantly increases neuronal survival (50). Additionally, wild-type α-syn has been reported to block rotenone- and maneb-induced ROS production and cell toxicity. In contrast, the mutated versions of α-syn, such as A30P, A53T, and E46K, aggravated ROS production induced by rotenone and maneb (51). Moreover, in one of the studies involving microglial cells, the administration of aggregated wild-type α-syn showed a dose-dependent increase in ROS production, while non-aggregated α-syn resulted in a slightly decreased ROS level (52). Thus, the ability of α-syn to modulate ROS appears to be dependent on the conformational form of the protein. In terms of JEV progression in neuronal cells, Exoα-syn was found to modulate oxidative stress inside the cell. Furthermore, it blocked the JEV-induced ROS generation during the late phase of JEV infection.

We investigated whether the anti-JEV activity exhibited by α-syn is mediated via modulation of ROS levels. The expression of anti-oxidative proteins like NQO1, SOSM1, and DJ-1, that are mainly involved in Nrf2-SQSTM1 pathway, was not altered with Exoα-syn protein treatment. However, there was upregulation of SOD1 with Exoα-syn treatment and conversely, endogenous α-syn silenced neuro2a cells had low SOD1 expression. This anti-oxidative mechanism of α-syn has been shown earlier, wherein SH-SY5Y cells expressing α-syn protein had increased SOD1 activity and significantly attenuated rotenone-induced cell apoptosis (53). SOD1 is known to convert the superoxide radicals arising from mitochondrial intermembrane space, cytosol, and peroxisome into hydrogen peroxide and molecular oxygen through redox reactions. Overall, our results indicate that α-syn, through SOD1 upregulation, contributes to a cellular environment that counteracts oxidative stress and restrains JEV propagation. This was supported by the data where inhibiting SOD1 activity using the compound LCS-1 led to an increased release of JEV particles outside the cells. We did not observe a significant difference in intracellular viral proteins with LCS-1 treatment, indicating the late response of SOD1 which may be during packaging or budding of viral particles outside cells. SOD1 is a known target of a plant lignan compound, arctigenin that shows anti-JEV activity via increasing SOD1 protein level inside cells (54). In JEV-infected *in vivo* and *in vitro* models, SOD1 is reported to be downregulated (54). Like SOD1, there are many other anti-oxidative proteins that are manipulated differentially by most of the *flaviviridae* members. JEV is reported to decrease ROS scavenging *via* downregulation of proteins like DJ-1 as well as Nrf2-mediated expression of SQSTM1 and thioredoxin (37, 55). Thus, there is a significant role of oxidative stress-induced ROS in shaping the cellular milieu to favor JEV propagation. This is further established by the data where administration of the oxidative stress inducer, hydrogen peroxide (H_2_O_2_), significantly increased viral replication within infected cells.

In conclusion, our study sheds light on the intricate interplay between oxidative stress, α-syn, and JEV infection. Apart from current study, α-syn is reported to modulate ER stress pathway (42). The levels of activating transcription factor 6 (Atf6), protein disulfide isomerase (PDI), and phosphorylated eIF2α proteins, all of which support viral infection in WNV were significantly increased in α-syn-knockout primary cortical neurons (42). Thus, there is possibility of more than one antiviral mechanism or pathway being involved in response to α-syn. It is noteworthy that we have not investigated the role of aggregated α-syn or Lewy bodies (LBs) in JEV infection. As maintaining the physiologic levels of α-syn in neurons is important for their survival, overexpression of α-syn for prolonged time may lead to its aggregation even if it is in response of viral infection. Therefore, the role of LBs and aggregation of α-syn in viral pathogenesis remains an important question and an area of active interest.

## CRediT authorship contribution statement

**Anjali Gupta:** Conceptualization, Data curation, Formal analysis, Investigation, Methodology, Visualization, Writing - original draft. **Anshuman Mohapatra**: Methodology, Formal analyses, Investigation, Writing-synthesis draft. **Ajanta Sharma**: Investigation, Resources. **Harpreet Kaur**: Resources. **Nitin Chaudhary**: Resources, Investigation, Methodology, Formal analyses, Supervision, Writing-review and editing. **Sachin Kumar**: Conceptualization, Funding acquisition, Investigation, Project administration, Resources, Supervision, Validation, Writing-review and editing.

## Conflict of interest

The authors declare no conflict of interest.

## Acknowledgments

Funding from Department of Health Research, Government of India (Grant No. NER/71/2020-ECD-I) is gratefully acknowledged.

---

**Supplementary Figure 1**. Characterization of purified Exoα-syn. The SEC profile of the anion exchange-purified αS (A). A 12% SDS-PAGE of Exoα-syn SEC fractions. The BioRad Precision Plus Protein™ ladder marked as M. The SEC fractions were loaded in L1-L8 (B). MALDI-TOF mass spectrometric analyses of Exoα-syn protein. Expected monoisotopic mass: 14,451.22 Da, observed m/z: 14,476.18. Doubly-charged species is seen at m/z 7,233.25 (C). DLS profile of the freshly prepared Exoα-syn protein (D).

**Supplementary Figure 2**. Characterization of liposomes (SUVs) and interaction with Exoα-syn protein. Graphs representing DLS size distribution profile of the plain and calcein-loaded SUVs (A). Circular dichroism spectra of Exoα-syn (2 μM) in phosphate buffer and in the presence of SUVs at protein:lipid ratio of 1:200 (B). Graphs representing fluorescence intensities of calcein-loaded SUVs in buffer, in the presence of Triton X-100 (1% v/v), and with Exoα-syn at 1:50 protein:lipid ratio (C).

## References

1. Murgod UA, Muthane UB, Ravi V, Radhesh S, Desai A. 2001. Persistent movement disorders following Japanese encephalitis. Neurology 57:2313–5.

2. Banerjee A, Tripathi A. 2019. Recent advances in understanding Japanese encephalitis. F1000Res 8.

3. Basumatary LJ, Raja D, Bhuyan D, Das M, Goswami M, Kayal AK. 2013. Clinical and radiological spectrum of Japanese encephalitis. J Neurol Sci 325:15–21.

4. Srivastava R, Kalita J, Khan MY, Gore MM, Bondre VP, Misra UK. 2013. Temporal changes of Japanese encephalitits virus in different brain regions of rat. Indian J Med Res 138:219–23.

5. Tadokoro K, Ohta Y, Sato K, Maeki T, Sasaki R, Takahashi Y, Shang J, Takemoto M, Hishikawa N, Yamashita T, Lim CK, Tajima S, Abe K. 2018. A Japanese Encephalitis Patient Presenting with Parkinsonism with Corresponding Laterality of Magnetic Resonance and Dopamine Transporter Imaging Findings. Intern Med 57:2243–2246.

6. Misra UK, Kalita J. 2002. Prognosis of Japanese encephalitis patients with dystonia compared to those with parkinsonian features only. Postgrad Med J 78:238–41.

7. DeMaagd G, Philip A. 2015. Parkinson’s Disease and Its Management: Part 1: Disease Entity, Risk Factors, Pathophysiology, Clinical Presentation, and Diagnosis. P T 40:504–32.

8. Jankovic J, Tan EK. 2020. Parkinson’s disease: etiopathogenesis and treatment. J Neurol Neurosurg Psychiatry 91:795–808.

9. Limphaibool N, Iwanowski P, Holstad MJV, Kobylarek D, Kozubski W. 2019. Infectious Etiologies of Parkinsonism: Pathomechanisms and Clinical Implications. Front Neurol 10:652.

10. Hopkins HK, Traverse EM, Barr KL. 2022. Viral Parkinsonism: An underdiagnosed neurological complication of Dengue virus infection. PLoS Negl Trop Dis 16:e0010118.

11. Koyuncu OO, Hogue IB, Enquist LW. 2013. Virus infections in the nervous system. Cell Host Microbe 13:379–93.

12. Burre J, Sharma M, Sudhof TC. 2018. Cell Biology and Pathophysiology of alpha-Synuclein. Cold Spring Harb Perspect Med 8.

13. Bussell R, Jr., Eliezer D. 2003. A structural and functional role for 11-mer repeats in alpha-synuclein and other exchangeable lipid binding proteins. J Mol Biol 329:763–78.

14. Liu C, Zhao Y, Xi H, Jiang J, Yu Y, Dong W. 2021. The Membrane Interaction of Alpha-Synuclein. Front Cell Neurosci 15:633727.

15. Pfefferkorn CM, Jiang Z, Lee JC. 2012. Biophysics of alpha-synuclein membrane interactions. Biochim Biophys Acta 1818:162–71.

16. Westphal CH, Chandra SS. 2013. Monomeric synucleins generate membrane curvature. J Biol Chem 288:1829–40.

17. Mizuno N, Varkey J, Kegulian NC, Hegde BG, Cheng N, Langen R, Steven AC. 2012. Remodeling of lipid vesicles into cylindrical micelles by alpha-synuclein in an extended alpha-helical conformation. J Biol Chem 287:29301–11.

18. Varkey J, Isas JM, Mizuno N, Jensen MB, Bhatia VK, Jao CC, Petrlova J, Voss JC, Stamou DG, Steven AC, Langen R. 2010. Membrane curvature induction and tubulation are common features of synucleins and apolipoproteins. J Biol Chem 285:32486–93.

19. Jenco JM, Rawlingson A, Daniels B, Morris AJ. 1998. Regulation of phospholipase D2: selective inhibition of mammalian phospholipase D isoenzymes by alpha- and beta-synucleins. Biochemistry 37:4901–9.

20. Gorbatyuk OS, Li S, Nguyen FN, Manfredsson FP, Kondrikova G, Sullivan LF, Meyers C, Chen W, Mandel RJ, Muzyczka N. 2010. alpha-Synuclein expression in rat substantia nigra suppresses phospholipase D2 toxicity and nigral neurodegeneration. Mol Ther 18:1758–68.

21. Oueslati A, Fournier M, Lashuel HA. 2010. Role of post-translational modifications in modulating the structure, function and toxicity of alpha-synuclein: implications for Parkinson’s disease pathogenesis and therapies. Prog Brain Res 183:115–45.

22. Stefanis L. 2012. alpha-Synuclein in Parkinson’s disease. Cold Spring Harb Perspect Med 2:a009399.

23. Bernal-Conde LD, Ramos-Acevedo R, Reyes-Hernandez MA, Balbuena-Olvera AJ, Morales-Moreno ID, Arguero-Sanchez R, Schule B, Guerra-Crespo M. 2019. Alpha-Synuclein Physiology and Pathology: A Perspective on Cellular Structures and Organelles. Front Neurosci 13:1399.

24. Irwin DJ, Lee VM, Trojanowski JQ. 2013. Parkinson’s disease dementia: convergence of alpha-synuclein, tau and amyloid-beta pathologies. Nat Rev Neurosci 14:626–36.

25. Mittal M, Siddiqui MR, Tran K, Reddy SP, Malik AB. 2014. Reactive oxygen species in inflammation and tissue injury. Antioxid Redox Signal 20:1126–67.

26. Schieber M, Chandel NS. 2014. ROS function in redox signaling and oxidative stress. Curr Biol 24:R453–62.

27. Heiden DL, Monogue B, Ali MDH, Beckham JD. 2023. A functional role for alpha-synuclein in neuroimmune responses. J Neuroimmunol 376:578047.

28. Roodveldt C, Labrador-Garrido A, Gonzalez-Rey E, Fernandez-Montesinos R, Caro M, Lachaud CC, Waudby CA, Delgado M, Dobson CM, Pozo D. 2010. Glial innate immunity generated by non-aggregated alpha-synuclein in mouse: differences between wild-type and Parkinson’s disease-linked mutants. PLoS One 5:e13481.

29. Somayaji M, Lanseur Z, Choi SJ, Sulzer D, Mosharov EV. 2021. Roles for alpha-Synuclein in Gene Expression. Genes (Basel) 12.

30. Schaser AJ, Osterberg VR, Dent SE, Stackhouse TL, Wakeham CM, Boutros SW, Weston LJ, Owen N, Weissman TA, Luna E, Raber J, Luk KC, McCullough AK, Woltjer RL, Unni VK. 2019. Alpha-synuclein is a DNA binding protein that modulates DNA repair with implications for Lewy body disorders. Sci Rep 9:10919.

31. Simanjuntak Y, Liang JJ, Lee YL, Lin YL. 2017. Japanese Encephalitis Virus Exploits Dopamine D2 Receptor-phospholipase C to Target Dopaminergic Human Neuronal Cells. Front Microbiol 8:651.

32. Kim SJ, Kim SY, Na YS, Lee HJ, Chung KC, Baik JH. 2006. Alpha-synuclein enhances dopamine D2 receptor signaling. Brain Res 1124:5–9.

33. Mohapatra A, Bohara VS, Kumar S, Chaudhary N. 2021. Polymyxin B accelerates the alpha-synuclein aggregation. Biophys Chem 277:106628.

34. Mohapatra A, Lokappa SB, Chaudhary N. 2020. Interaction of cavin-1/PTRF leucine zipper domain 2 and its congenital generalized lipodystrophy mutant with model membranes. Biochem Biophys Res Commun 521:732–738.

35. Chaudhary N, Nagaraj R. 2009. Hen lysozyme amyloid fibrils induce aggregation of erythrocytes and lipid vesicles. Mol Cell Biochem 328:209–15.

36. Furlong RA, Narain Y, Rankin J, Wyttenbach A, Rubinsztein DC. 2000. Alpha-synuclein overexpression promotes aggregation of mutant huntingtin. Biochem J 346 Pt 3:577–81.

37. Gupta A, Gawandi S, Vandna, Yadav I, Mohan H, Desai VG, Kumar S. 2022. Analysis of fluoro based pyrazole analogues as a potential therapeutics candidate against Japanese encephalitis virus infection. Virus Res 323:198955.

38. Mohapatra A, Hans A, Chaudhary N. 2023. Interfacial properties of alpha-synuclein’s Parkinsonian variants. Biophys Chem 297:107006.

39. Mohapatra A, Chaudhary N. 2021. N-terminal acetylation does not alter alpha-synuclein’s interfacial properties. Int J Biol Macromol 174:69–76.

40. Al-Obaidi MMJ, Bahadoran A, Wang SM, Manikam R, Raju CS, Sekaran SD. 2018. Disruption of the blood brain barrier is vital property of neurotropic viral infection of the central nervous system. Acta Virol 62:16–27.

41. Pardridge WM. 2012. Drug transport across the blood-brain barrier. J Cereb Blood Flow Metab 32:1959–72.

42. Beatman EL, Massey A, Shives KD, Burrack KS, Chamanian M, Morrison TE, Beckham JD. 2015. Alpha-Synuclein Expression Restricts RNA Viral Infections in the Brain. J Virol 90:2767–82.

43. Khanlou N, Moore DJ, Chana G, Cherner M, Lazzaretto D, Dawes S, Grant I, Masliah E, Everall IP, Group H. 2009. Increased frequency of alpha-synuclein in the substantia nigra in human immunodeficiency virus infection. J Neurovirol 15:131–8.

44. Wewer CR, Khandelia H. 2018. Different footprints of the Zika and dengue surface proteins on viral membranes. Soft Matter 14:5615–5621.

45. Angelova PR, Horrocks MH, Klenerman D, Gandhi S, Abramov AY, Shchepinov MS. 2015. Lipid peroxidation is essential for alpha-synuclein-induced cell death. J Neurochem 133:582–9.

46. Harischandra DS, Jin H, Anantharam V, Kanthasamy A, Kanthasamy AG. 2015. alpha-Synuclein protects against manganese neurotoxic insult during the early stages of exposure in a dopaminergic cell model of Parkinson’s disease. Toxicol Sci 143:454–68.

47. Zhu M, Qin ZJ, Hu D, Munishkina LA, Fink AL. 2006. Alpha-synuclein can function as an antioxidant preventing oxidation of unsaturated lipid in vesicles. Biochemistry 45:8135–42.

48. Kim JY, Jeon BS, Kim HJ, Ahn TB. 2013. Nanomolar concentration of alpha-synuclein enhances dopaminergic neuronal survival via Akt pathway. Neural Regen Res 8:3269–74.

49. Kaul S, Anantharam V, Kanthasamy A, Kanthasamy AG. 2005. Wild-type alpha-synuclein interacts with pro-apoptotic proteins PKCdelta and BAD to protect dopaminergic neuronal cells against MPP+-induced apoptotic cell death. Brain Res Mol Brain Res 139:137–52.

50. Xu J, Kao SY, Lee FJ, Song W, Jin LW, Yankner BA. 2002. Dopamine-dependent neurotoxicity of alpha-synuclein: a mechanism for selective neurodegeneration in Parkinson disease. Nat Med 8:600–6.

51. Choong CJ, Say YH. 2011. Neuroprotection of alpha-synuclein under acute and chronic rotenone and maneb treatment is abolished by its familial Parkinson’s disease mutations A30P, A53T and E46K. Neurotoxicology 32:857–63.

52. Thomas MP, Chartrand K, Reynolds A, Vitvitsky V, Banerjee R, Gendelman HE. 2007. Ion channel blockade attenuates aggregated alpha synuclein induction of microglial reactive oxygen species: relevance for the pathogenesis of Parkinson’s disease. J Neurochem 100:503–19.

53. Liu YY, Zhao HY, Zhao CL, Duan CL, Lu LL, Yang H. 2006. [Overexpression of alpha-synuclein in SH-SY5Y cells partially protected against oxidative stress induced by rotenone]. Sheng Li Xue Bao 58:421–8.

54. Swarup V, Ghosh J, Mishra MK, Basu A. 2008. Novel strategy for treatment of Japanese encephalitis using arctigenin, a plant lignan. J Antimicrob Chemother 61:679–88.

55. Yang TC, Lai CC, Shiu SL, Chuang PH, Tzou BC, Lin YY, Tsai FJ, Lin CW. 2010. Japanese encephalitis virus down-regulates thioredoxin and induces ROS-mediated ASK1-ERK/p38 MAPK activation in human promonocyte cells. Microbes Infect 12:643–51.

